# FerriTag: A Genetically-Encoded Inducible Tag for Correlative Light-Electron Microscopy

**DOI:** 10.1101/095208

**Authors:** Nicholas I. Clarke, Stephen J. Royle

## Abstract

A current challenge is to develop tags to precisely visualize proteins in cells by light and electron microscopy. Here, we introduce FerriTag, a genetically-encoded chemically-inducible tag for correlative light-electron microscopy (CLEM). FerriTag is a fluorescent recombinant electron-dense ferritin particle that can be attached to a protein-of-interest using rapamycin-induced heterodimerization. We demonstrate the utility of FerriTag for CLEM by labeling proteins associated with various intracellular structures including mitochondria, plasma membrane, and clathrin-coated pits and vesicles. FerriTagging has a high signal-to-noise ratio and a labeling resolution of 10 ± 5 nm. We demonstrate how FerriTagging allows nanoscale mapping of protein location relative to a subcellular structure, and use it to detail the distribution of huntingtin-interacting protein 1 related (HIP1R) in clathrin-coated pits.

## Introduction

To understand cell biology, we must explore subcellular organization in 3D and locate proteins at high resolution. Correlative light-electron microscopy (CLEM) is a powerful technique to do this, since we can combine the specificity and dynamics of fluorescence light microscopy (LM) with the high resolution and cellular context of electron microscopy (EM). A current challenge is to develop tools that allow us to track intracellular events using CLEM. Immunogold labeling has long been used for this purpose, however, it is particularly invasive and its applications are limited^1,2^. More recently, attention has turned to genetically-encoded tags for CLEM.

The ideal tag for CLEM should meet the following criteria: (1) fluorescent and electron dense so that it can be visualized by LM and EM, (2) the electron density should be tightly focused and provide good signal-to-noise ratio (SNR) so the tag itself is easily distinguishable from background by EM, (3) genetically encoded so that the cell can be processed in its native state without the need for permeabilization, and (4) non-toxic and non-disruptive so as not to interfere with normal cellular function.

Existing tags do not meet all of these criteria. A popular approach has been to use diaminobenzidine (DAB) to form an electron-dense precipitate either by enzymatic-based polymerization using peroxidase^3,4^ or singlet oxygen-based polymerization during photo-oxidation^5,6^. Although these tags allow CLEM, they result in low labeling resolution by EM due to the diffuse nature of the precipitate. Therefore, only proteins situated inside organelles, or those discretely localized at high densities can be successfully visualized.

The search for an ideal tag for CLEM has continued towards metal ligand-based tags combined with fluorescent proteins. Metal clusters contain elements that are able to scatter electrons and, if tightly focused, should improve resolution and be readily distinguishable from background. Two such tags are concatenated metallothionein^7,8^ and bacterioferritin^9^, however each have significant drawbacks in their current form. Use of metallothioneins is limited to high abundance proteins and only minimal ultrastructural information is currently possible^7^. Tagging with bacterioferritin is technically demanding, limited to bacteria, and can lead to aggregation and mislocalization of the target^9^.

Human ferritin is a complex of 24 polypeptide subunits of light (FTL) and/or heavy (FTH) chains that form a spherical protein shell with internal and external diameters of approximately 7 and 12 nm respectively. Under iron-rich conditions it is able to store iron (<4300 Fe(III) atoms) as a mineral core and is therefore easily distinguishable by EM^10^. The electron density of Ferritin has been exploited for immunoEM for decades^11,12^ and we hypothesized that it could be used as an ideal CLEM tag with some modifications.

Here we introduce FerriTag, a genetically-encoded inducible tag for CLEM. FerriTag has been engineered to build an electron-dense recombinant ferritin particle that can label any protein-of-interest using rapamycin-induced heterodimerization. We demonstrate how FerriTag can be used to acutely label single proteins at nanometer resolution and highlight how this new technology will be useful in answering key questions in cell biology.

## Results

### Design and implimentation of FerriTagging

FerriTagging involves the creation of a ferritin particle (FerriTag) which can be inducibly attached to a protein-of-interest using the FKBP-rapamycin-FRB heterodimerization system (Figure 1A). To do this, FerriTag is co-expressed with the protein-of-interest which is fused to FKBP-GFP. FerriTag is untagged FTL and FRB-mCherry-FTH1 transfected at a ratio of 4:1. In the absence of rapamycin, the target protein performs its normal cellular function. When rapamycin is added, it induces the heterodimerization of FKBP and FRB domains resulting in the target protein becoming FerriTagged (Figure 1A).

**Figure 1:**
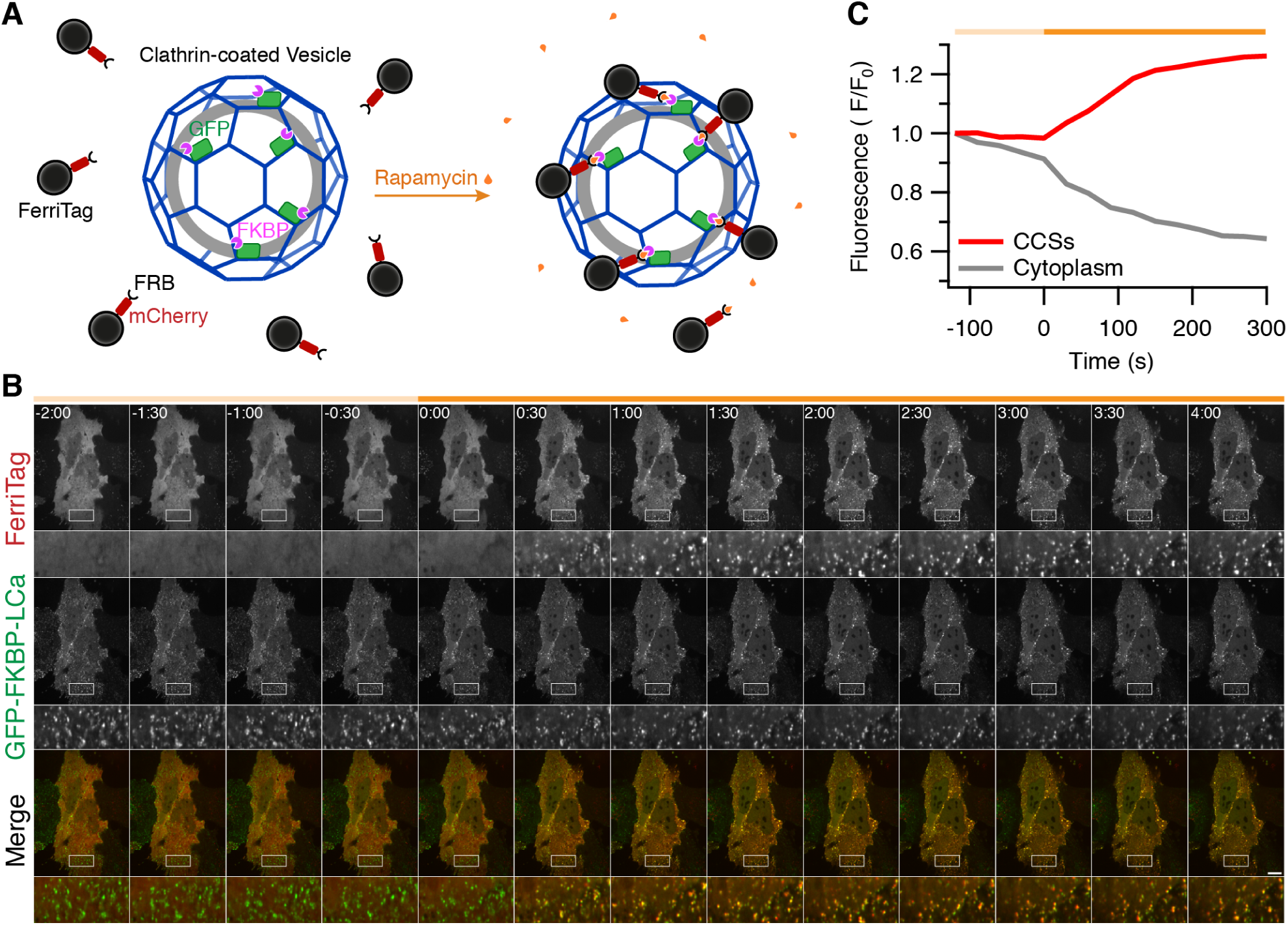
Design and implimentation of FerriTagging. **(A)** Schematic diagram of FerriTagging clathrin light chain. Simultaneous expression of FerriTag and GFP-FKBP-tagged clathrin light chain. Addition of rapamycin induces the heterodimerization of FKBP and FRB domains resulting in FerriTagging of clathrin in a clathrin-coated vesicle. **(B)** Stills from live-cell imaging of FerriTagging clathrin light chain. Rapamycin (200 nmoll^−1^) was added at time point zero, as indicated by the orange bar. Specific, rapid labeling of clathrin by FerriTag can be observed in <30s. Time, min: sec, scale bar, 10 μm. Zooms show 4x expansion. See **Supplementary Video 1**. **(C)** Quantification of FerriTag (mCherry) fluorescence in clathrin-coated structures (CCSs, red) and in the cytoplasm (gray) over time during the experiment shown in B.

Our initial attempt to use Ferritin as a CLEM tag involved the direct fusion of FTH1 to a protein-of-interest, as described previously^9^. Mammalian cells expressing mCherry-FTH1 fused to the mitochondrial targeting sequence of Tom70p were cultured under iron-rich conditions and then visualized by fluorescence microscopy (**Supplementary Figure 1**). It was clear by LM that this direct fusion resulted in aggregation and mislocalization of mitochondria, most likely due to the multivalent nature of the ferritin molecule. This observation drove us to engineer a novel ferritin tag that could form a ferritin particle independently and then be inducibly added to a protein-of-interest to avoid such aggregation issues. The dilution of FRB-mCherry-tagged FTH1 subunits with untagged FTL subunits is also an important step because aggregation was observed when FRB-mCherry-FTH1 was expressed alone (**Supplementary Figure 1**). The outcome of this diluted-tagged version of ferritin, was no aggregation or mislocalization issues before or after the addition of rapamycin (**Supplementary Figure 1**).

Our first target protein for FerriTagging was clathrin. HeLa cells expressing FerriTag and clathrin light chain a (LCa) tagged with GFP and FKBP were imaged by fluorescence microscopy (Figure 1B). These experiments show that, after the addition of rapamycin (200 nmoll^−1^), FerriTagging of clathrin-coated structures occurs within seconds (Figure 1C). The diffuse FerriTag signal can be seen to specifically decorate clathrin-coated structures throughout the entire cell within minutes (Figure 1B **and Supplementary Video 1**). With this approach working, we next wanted to observe FerriTagging by EM.

### Visualizing FerriTagged proteins by electron microscopy

In order to assess whether or not FerriTagging works at the EM level, we used correlative light-electron microscopy (CLEM). Our EM protocol was optimized so that we could easily distinguish FerriTag from background and be able to see cellular ultrastructure (Figure 2A, see Methods). We initially FerriTagged two different proteins in disparate locations to assess this optimized protocol for CLEM: 1) monoamine oxidase (MAO) an outer mitochondrial membrane protein, and 2) clathrin light chain a (LCa), part of the clathrin triskelion (Figure 2B). Following the addition of rapamycin, FerriTag was confirmed by fluorescence microscopy to specifically label each protein-of-interest rapidly. After fixation and processing, ultrathin resin sections taken from the same cell were imaged by EM. In each case, electron-dense particles of approximately 7 nm diameter could be easily distinguished from background, and the ultrastructure of the cell was visible. We were able to locate particles specifically in the vicinity of mitochondria (in the case of MAO) or CCSs (in the case of LCa) where the protein-of-interest is localized (Figure 2B). Labeling was sparse in ultrathin 70 nm sections. Particles observed by EM most likely correspond to FerriTag because first, no labelling was seen at CCSs in cells where MAO was FerriTagged, and mitochondria were not tagged in cells where clathrin was FerriTagged, second, no particles were observed next to the appropriate organelle in cells where no rapamycin has been added, and nor when third, FerriTagging was done in cells with no FeSO_4_ loading (**Supplementary Figure 2**). From these experiments we conclude that FerriTag works to specifically label proteins-of-interest at the ultrastructural level.

**Figure 2:**
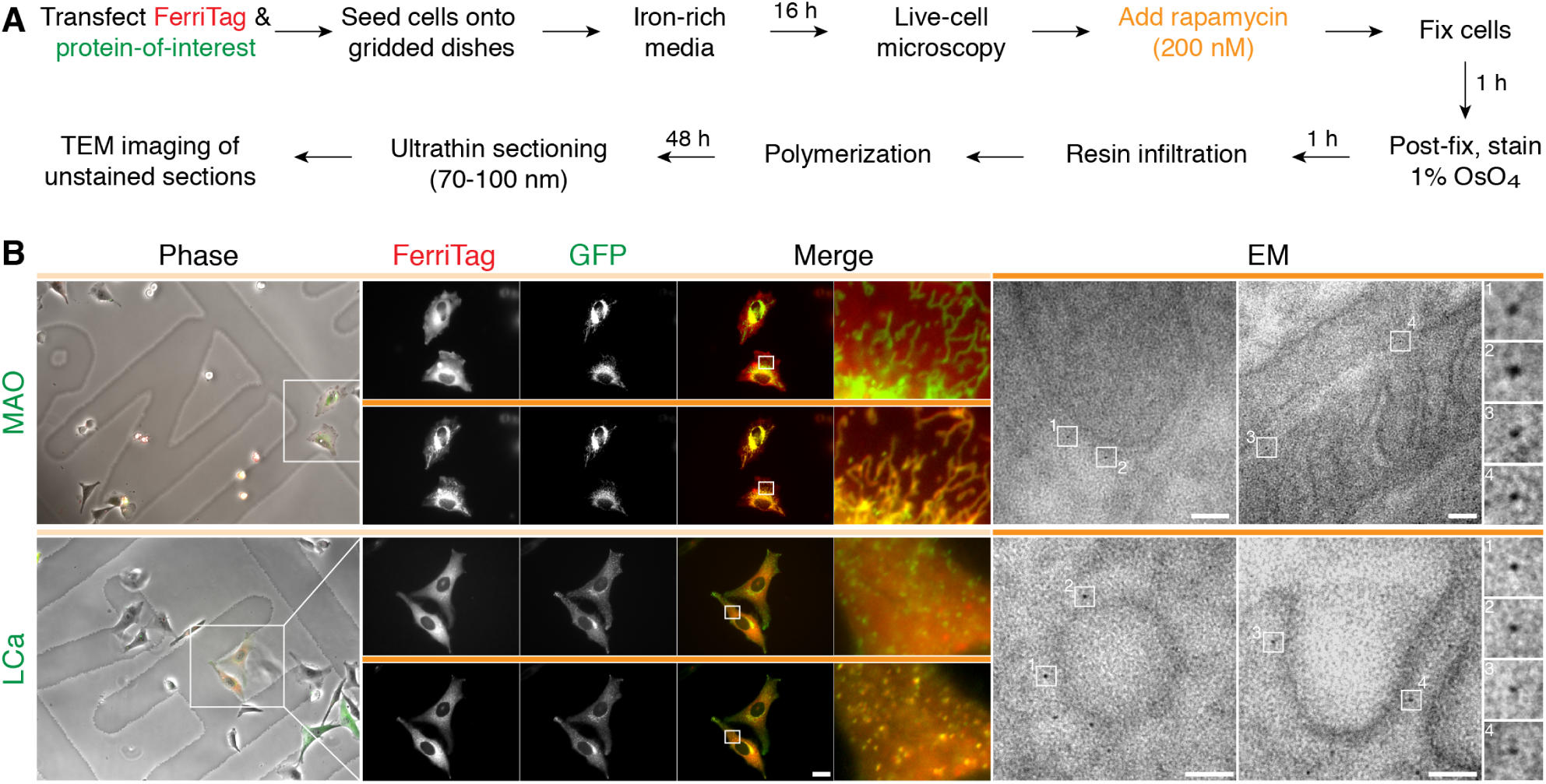
Visualizing FerriTagged proteins by LM and EM. **(A)** Overview of sample preparation steps for CLEM using FerriTag. **(B)** Light and electron micrographs of HeLa cells co-expressing FerriTag with either GFP-FKBP-Myc-MAO or GFP-FKBP-LCa. Locator images (left) ensure the same cell can be followed throughout the CLEM workflow. Live cell imaging of FerriTagging (middle), cells were treated with rapamycin (200 nmoll^−1^) as indicated by the filled orange bar. The last frame is shown before the cell was fixed and processed for CLEM as described in A. High-resolution TEM (right) of sections taken from the same cell show dense FerriTag particles specifically labeling the GFP-FKBP-tagged protein-of-interest in each CLEM experiment. Two particles per micrograph are shown expanded to the right. Light microscopy scale bar 10 μm and zoom ×12. Electron micrograph scale bar 50 nm and zoom ×7.25.

### Determining the labeling resolution of FerriTag

What is the labeling resolution of FerriTagging? Labeling resolution refers to the accuracy with which the detected position of the particle corresponds to the location of the labeled protein^2^. The maximum possible distance that the FerriTag particle could be away from a target protein is 22 nm (Figure 3A). However, FerriTag may adopt a more compact conformation or pose resulting in shorter observed lengths (see Supplementary Information). To determine the labeling resolution directly, we FerriTagged the transmembrane protein CD8*α* and processed samples for EM (Figure 3B. The perpendicular distance from the center of the FerriTag particle to plasma membrane was measured (median = 9.5 nm, N = 458 particles, Figure 3C). To interpret the shape of this distribution, we carried out computer simulations that modeled the detection of particles in EM sections (see Methods and Supplementary Information). These simulations indicate that FerriTag can exist in a number of length states from 7 to 18 nm. The distribution has a broad spread (FWHM ≈ 10 nm), which means that on average the particle will be 10 ± 5 nm away from the target protein. This resolution exceeds that of traditional immunogold labeling which has a labeling resolution of 18 nm in pre-embedding or 21 nm on-section^2^. This dataset also allowed us to directly determine the SNR for FerriTagging (SNR = 12 ± 0.2, mean ± s.e.m., see Methods and Supplementary Information, Figure 3D).

**Figure 3:**
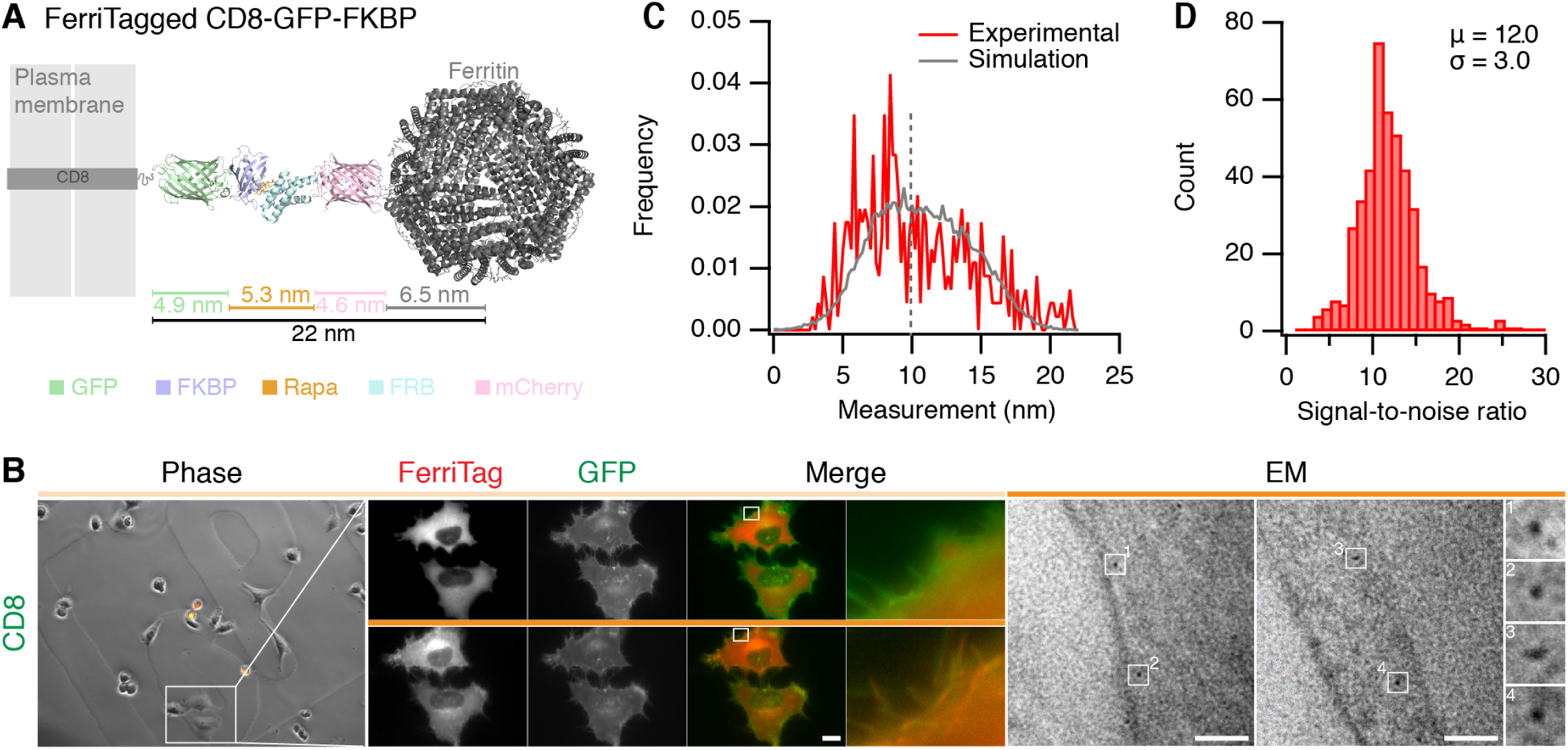
Labeling resolution of FerriTagging is approximately 10 nm. **(A)** Schematic diagram to show the experimental setup to measure labeling resolution of FerriTag. FerriTagging of CD8-GFP-FKBP is shown relative to the plasma membrane. Protein domains are organized co-linearly, giving a total maximum length of 22 nm from the edge of the plasma membrane to the center of the ferritin particle. **(B)** FerriTagging CD8-GFP-FKBP. Locator images (left), live cell imaging of FerriTagging (middle), and TEM images (right). Cells were treated with rapamycin (200 nmoll^−1^) as indicated by the filled orange bar. Light microscopy scale bar 10 μm and zoom ×12. Electron micrograph scale bar 50 nm and zoom ×7.25. **(C)** Histogram of experimental observations (red) with a density function of simulated values overlaid (gray). Dotted line indicates the median 9.8 nm (*N_particle_* = 458). **(D)** Histogram of signal-to-noise ratio (SNR) measured from this dataset (*N_particle_* = 441).

### Nanoscale mapping of HIP1R shows an even distribution at clathrin-coated pits

Having developed FerriTagging, we next wanted to carry out contextual nanoscale mapping of a protein-of-interest to answer a cell biological question. Huntingtin-interacting protein 1 related (HIP1R), the human homolog of yeast Sla2p, can bind clathrin light chain, actin and membranes and is proposed to serve as a link between the clathrin machinery to the actin cytoskeleton^13–16^. Current models of actin function during CME in human cells constrain the distribution of actin to an accumulation at the neck of deeply invaginated CCPs, and HIP1R is thought to have a corresponding distribution at the circumference of CCPs^17–19^. To test this model wanted to determine the nanoscale distribution of HIP1R at clathrin-coated pits using FerriTagging. HeLa cells expressing HIP1R-GFP-FKBP and FerriTag were processed through our CLEM workflow and high resolution electron micrographs of FerriTagged HIP1R on clathrin-coated pits were acquired (Figure 4A). We collected images of clathrin-coated pits and segmented the plasma membrane profile and position of FerriTag particles in each (Figure 4B, see Methods). Using spatial averaging, we plotted the distribution of all particles symmetrically about an idealized pit profile (Figure 4C). These data revealed that the distribution of HIP1R is homogenous over the entire crown of the CCP. Moreover, by defining the edges of the CCP we could map FerriTagged HIP1R *outside* of the CCP, i.e. in adjacent areas of plasma membrane. This analysis revealed lower density, but significant, labeling of HIP1R outside the CCP (Figure 4C,D). Interestingly, the distance from particles to the plasma membrane was greater for FerriTagged HIP1R inside the pit versus outside, a difference of 10 nm on average (Figure 4E). These observations suggest that the distribution of HIP1R is not constrained to the circumference of CCPs, but is homogenously distributed over the clathrin coat and in surrounding, uncoated areas of plasma membrane.

**Figure 4:**
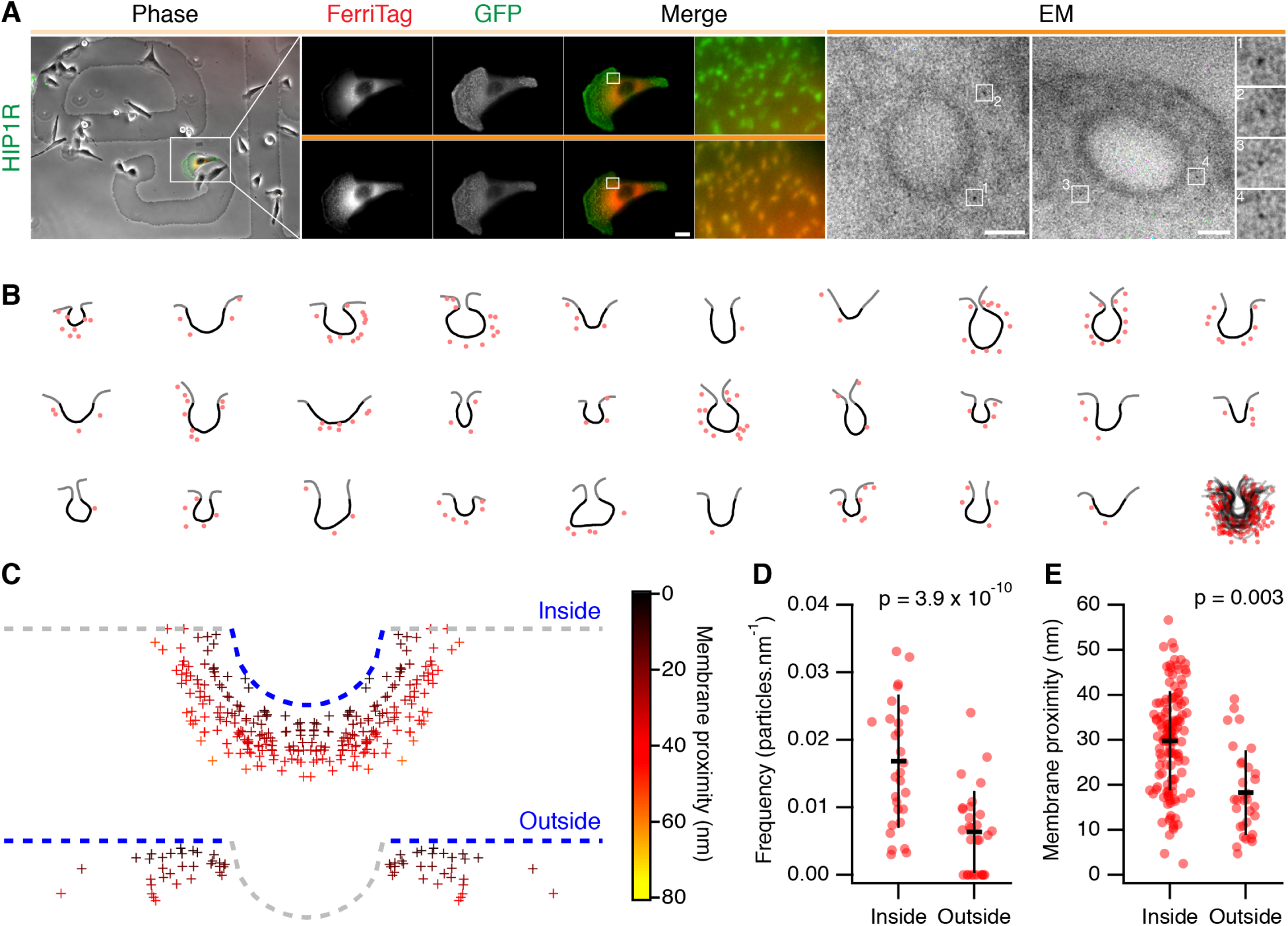
Nanoscale mapping of HIP1R location in the vicinity of clathrin-coated pits. **(A)** FerriTagging HIP1R-GFP-FKBP. Locator images (left), live cell imaging of FerriTagging (middle), and TEM images (right). Cells were treated with rapamycin (200 nmoll^−1^) as indicated by the filled orange bar. Light microscopy scale bar 10 μm and zoom ×12. Electron micrograph scale bar 50 nm and zoom ×7.25. **(B)** Manually segmented membrane profiles (Black-inside, Gray-outside) and FerriTag particles (red) from HIP1R FerriTag electron micrographs. **(C)** Spatially averaged representation of HIP1R bound to an idealized clathrin-coated pit determined by position of FerriTag particles in TEM micrographs. FerriTag locations are marked by crosses, color-coded by the distance to the plasma membane. Note, that the distribution is symmetrically presented, i.e. each particle appears twice. **(D-E)** Scatter dot plots of the frequency of FerriTagged-HIP1R (D) and the proximity of FerriTagged-HIP1R to the plasma membrane (E). Particles inside or outside of the pit are plotted. Bars show mean ± SD.

## Discussion

In this paper we described FerriTag, a genetically-encoded chemically-inducible tag for CLEM that can be used to acutely label proteins in mammalian cells. The fluorescence and electron density of FerriTag allows proteins to be tracked by fluorescence microscopy in live cells and then visualized at the nanoscale by EM.

FerriTag meets all four criteria for an ideal CLEM tag: 1) fluorescent and electron dense, 2) the electron density should be tightly focused, 3) genetically encoded, and 4) non-disruptive. Currently, the most widely used CLEM tags rely on the production of an electron dense cloud of precipitate that precludes precise localization of target proteins. Due to the tightly focused electron density and good SNR of ferritin, FerriTag can be used for nanoscale mapping of protein location. There have been previous attempts to use metal ligand-based tags to do this^7–9^, however there are significant limitations to these methods which has prevented their wide application.

The FerriTag protocol outlined here is simple and robust, with the potential to be used for any protein-of-interest which can be fused to FKBP. Ultrastructure is well preserved due to absence of detergent and it will also possible to incorporate high-pressure freezing into the protocol to better preserve ultrastructure^20^, something that is not currently possible with DAB-based genetically encoded tags. Potential future innovations of FerriTagging include: 1) alternative CLEM protocols that allow for ultra-precise correlated single spot fluorescence localization with FerriTag^21–23^, 2) improving staining to enhance the visualization of ultrastructure, 3) combining FerriTag with other tagging methods, perhaps with two color EM ^24^, allowing for multicolor EM, and 4) harnessing the magnetic properties of FerriTag for use as a purification tag, or perhaps for direct magnetic manipulation of proteins in living cells.

The SNR of CLEM tags imaged by EM is complex. We need sufficient contrast to be able to achieve contextual nanoscale mapping, but this necessarily reduces the SNR. Equally, thicker sections should allow greater sampling of particles, yet increase the noise and complicate detection. The SNR of FerriTagging (12:1) is good, but it may be possible to increase this. One current limitation of our method is that it might not be possible to tag proteins which are located inside organelles, since FerriTag must be able to access the FKBP for FerriTagging to occur. However, this can be easily assessed by light microscopy, before any samples are processed for EM.

Contextual nanoscale mapping of proteins allows investigators to detail the fine distribution of a protein-of-interest in the context of subcellular ultrastructure. In this paper, we used FerriTagging to do contextual nanoscale mapping of HIP1R in HeLa cells. Previous evidence from immunogold deep-etch EM work suggested that HIP1R is localized at the circumference of CCPs, where actin is predicted to accumulate to generate a propulsive force for vesicle production^17–19^. Instead we found that HIP1R is localized throughout the CCP and even in surrounding areas of uncoated membrane, albeit at lower density. One current model is that the N-terminal ANTH domain of HIP1R associates with the plasma membrane of the CCP, the middle domain binds the clathrin coat, and the C-terminal region THATCH domain binds actin^14^. Using FerriTagging, we found that HIP1R outside of the CCP was closer to the plasma membrane than HIP1R within the pit, a difference of ≈10 nm. This difference could be due to HIP1R being directly associated with the plasma membrane outside of the CCP and then within the pit, HIP1R is located on the outside face of the clathrin coat, not bound to the plasma membrane. Alternatively, HIP1R could be bound to the plasma membrane in both locations but adopts a compact conformation outside the CCP and an extended conformation on the inside.

The labeling resolution of FerriTag is 10 ± 5 nm, which exceeds the predicted resolution provided by standard immunogold labeling by either pre-embedding or on-section^2^. Furthermore, FerriTag exceeds the resolution of currently available super-resolution light microscopy methods, with the added benefit that cellular context can be observed in relation to nanoscale localization of the protein-of-interest^23,25^.

These are exciting times for exploration of the subcellular world and for investigations into protein function in cells at the nanoscale. We hope FerriTag will be widely adopted as a discovery tool on these expeditions.

## Methods

### Molecular biology

To make FRB-mCherry-FTH1, human ferritin heavy polypeptide 1 (IMAGE clone: 3459353) was amplified by PCR and inserted into pFRB-mCherry-C1 via *XhoI-Eco*RI. FKBP-GFP-Myc-MAO was a kind gift from Sean Munro (MRC-LMB, Cambridge)^26^. For expression of FTL only, human ferritin light polypeptide cDNA (IMAGE clone: 2905327) was amplified by PCR and inserted into pEGFP-C1, removing EGFP, via *AgeI-Xho*I. GFP-FKBP-LCa was available from previous work^27^. To make CD8-GFP-FKBP, CD8α was amplified by PCR and inserted into pEGFP-FKBP-N1 via *NheI-AgeI*. To make HIP1R-GFP-FKBP, HIP1R (Addgene plasmid 27700) was amplified by PCR and inserted into pEGFP-FKBP-N1 via *XhoI-AgeI*. To make pMito-mCherry-FTH1, FTH1 was inserted into pMito-mCherry-FRB (from earlier work^27^) via *Bsr*G1-*Xba*I.

### Cell biology

HeLa cells were cultured in Dulbecco’s Modified Eagle Medium (Invitrogen) supplemented with 10% fetal bovine serum and 100 U/ml penicillin/ streptomycin at 37°C and 5% CO_2_. Cells were transfected with a total of 1.5 μg DNA (for 3.5 cm dishes) using Genejuice (Novagen) following manufacturer’s instructions. The total amount of DNA for each plasmid transfected in FerriTag experiments, unless otherwise specified, was 750 ng for GFP-FKBP tagged protein of interest, 600 ng for FTL only vector and 150 ng for FRB-mcherry-FTH1. Cells were imaged or fixed 2d after transfection. Cells grown in iron-rich conditions were supplemented with FeSO_4_ to a final concentration of 1 mmo||^−1^ in growth media, 16 h prior to imaging. Culturing cells in 1 mmo||^−1^ FeSO_4_ for this period of time has minimal effect on cell viability and does not alter cellular ultrastructure^28^.

### Light Microscopy

Live cell imaging of FerriTag kinetics was performed on a spinning disc confocal microscope (Ultraview Vox, Perkin Elmer) with a 100× 1.4 NA oil-immersion objective at 37 °C. Cells were cultured in glass-bottom fluorodishes (WPI) and kept in Leibovitz L-15 CO_2_-independent medium supplemented with 10% FBS during imaging. Images were captured using an ORCA-R2 digital CCD camera (Hamamatsu) following excitation with 488 nm and 561 nm lasers.

Fixed cell experiments were performed in transiently transfected cells attached to cover slips and fixed with 3% paraformaldehyde, 4% sucrose in PBS at 37°C. Cells were then washed in PBS before being mounted in Mowiol containing DAPI. Imaging was performed on a Nikon Ti epiflorescence microscope with standard filtersets, equipped with a heated environmental chamber (OKOlab) and CoolSnap MYO camera (Photometrics) using NIS elements AR software. Where applicable, rapamycin (Alfa Aesar) was added by flowing in a concentrated solution in growth media at 37°C to a final concentration of 200 nmoll^−1^.

### Correlative light electron microscopy

Following transfection, cells were plated onto gridded glass MatTek dishes (P35G-1.5-14-CGRD, MatTek Corporation, Ashland, MA, USA). Light microscopy was performed as described above. Cells were kept at 37°C in Leibovitz L-15 CO_2_-independent medium supplemented with 10% FBS during imaging. Transiently expressing cells were located and the photo-etched grid coordinate containing the cell of interest was recorded using brightfield illumination at 20x for future reference. The same cell was then relocated and fluorescent live cell imaging was then acquired at 100x. During imaging, rapamycin was added and once sufficient labeling had been achieved, cells were immediately fixed in 3% glutaraldehyde, 0.5% paraformaldehyde in 0.05 moll^−1^ phosphate buffer pH 7.4 for 1h. Following fixation, free aldehydes were quenched in 50 mmo||^−1^ glycine solution and cells washed several times in 0.05 mo||^−1^ phosphate buffer. Cells were post-fixed in 1% osmium tetroxide (Agar) for 1 h, washed in distilled water and then dehydrated through an ascending series of ethanol prior to infiltration with epoxy resin (TAAB) and polymerization at 60°C. This gave sufficient contrast without the need for post-staining. Coverslips attached to the polymerized resin block were removed by briefly plunging into liquid nitrogen. The cell of interest was then located by correlating grid coordinates imprinted on the resin block with previously acquired brightfield images. The resin around the cell of interest was then trimmed away using a glass knife. Serial, ultrathin sections of 70 nm were then taken using a diamond knife on an EM UC7 (Leica Microsystems) and collected on uncoated hexagonal 100 mesh grids (EM resolutions). Electron micrographs were recorded using a JEOL 1400 TEM operating at 100 kV using iTEM software.

### Image analysis and computer simulation

To measure labeling resolution, the location of the Ferritin particle and the plasma membrane were recorded using `Fiji`. These measurements were used to construct the histogram and this probability density function was used for comparison during computer modeling (see Supplementary Information). Briefly, FerriTag was modeled as a ball-and-chain with a variety of length states from 6.5 to 22 nm fixed at the origin on an *xy* plane at *z* = 0. The simulation assumed that FerriTag can be posed in a number of conformations such that the centre of Ferritin corresponds to the spherical coordinate triplet,

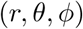

giving the coordinates,

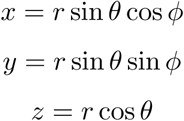

where *r* ∈ (6.5, 22], 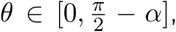 and *ϕ* ∈ [0, 2π]. Exclusion of *α* rad from the upper limit of *θ* is necessary because the outer edge of Ferritin may only touch the *xy* plane at *z* = 0, but not pass through it. Accordingly, *α* is calculated by

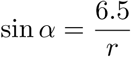

The simulation included a final parameter, noise (*σ* = 1.5) to allow for measurement error. A simulation where an equal number of a range of length states from 7 to 18 nm in 1nm steps matched the observed data well (see Supplementary Information for further details.

To determine the signal-to-noise ratio (SNR) of FerriTagging, we used the same dataset implementing a simple image analysis pipeline. The *xy* coordinates of each particle were used to excise a 11×11 pixel box centered on the particle. This image clip was fitted with a 2D Gaussian function,

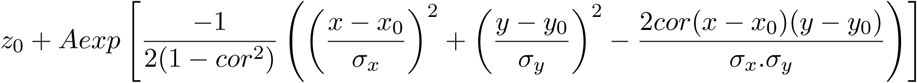

where the cross-correlation term, *cor* ∈ [−1,1] (see Supplementary Information). The image was inverted and the peak location (*x*_0_,*y*_0_) was used to center a 3×3 pixel ROI to calculate *μ_signal_*. A 50×50 pixel region in the cell, avoiding any extracellular areas or grid bars, was used to calculate *σ_backgrourad_*, to give

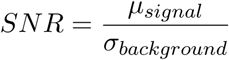

Code for this analysis is available at https://github.com/quantixed/FerriTag.

For mapping of HIP1R FerriTagging, electron micrographs in TIFF format were imported into IMOD and the plasma membrane and location of electron dense particles was manually segmented. The coordinates corresponding to contours and objects were fed into IgorPro 7.01 using the output from model2point. Custom-written procedures (available at https://github.com/quantixed/FerriTag) processed these data. First, the coordinates were scaled from pixels to real-world values, and the closest distance (proximity) to the plasma membrane was recorded. Next, the beginning and end of the pit were defined manually in a graphical user interface. The contour length of the pit was determined and the contour length between the start of the pit and the point of closest approach for each FerriTag particle was calculated. The ratio of these two lengths allowed us to plot out a spatially normalized view of the labeling locations. Particles that were closest to the membrane outside of the pit were plotted separately.

## Manuscript information

### Supplementary Information

1. Supplementary Information document PDF containing supplementary results, 7 supplementary figures, and a description of computer modeling
2. Supplementary Video 1 - FerriTagging clathrin in HeLa cells Example confocal live cell imaging of rapamycin (200 nM) application to HeLa cells expressing GFP-FKBP-LCa (green, left), FerriTag (red, middle), merge is shown to the right. See Figure 1. Imaging rate, 10 s per frame (green), 30 s per frame (red). Video speed, 10 fps.

## Acknowledgments

We would like to thank Natalie Allcock, Faye Nixon and Ian Prior for technical help. We also thank Sean Munro for reagents, and colleagues in CMCB for critical discussion throughout the project, particularly Rob Cross for reading our manuscript critically. This work was supported by a Senior Cancer Research UK Fellowship (C25425/A15182) to SJR and a Cancer Research UK Studentship (C25425/A16141) to NIC.

## Author contributions

NIC: did all experimental work, analyzed data, and wrote the paper, SJR: analyzed data, wrote Igor code, and wrote the paper.

